# Potential for *Methanosarcina* to contribute to uranium reduction during acetate-promoted groundwater bioremediation

**DOI:** 10.1101/202242

**Authors:** Dawn E Holmes, Roberto Orelana, Ludovic Giloteaux, Li-Ying Wang, Pravin Shrestha, Kenneth Williams, Derek R Lovley, Amelia-Elena Rotaru

## Abstract

Previous studies of *in situ* bioremediation of uranium-contaminated groundwater with acetate injections have focused on the role of *Geobacter* species in U(VI) reduction because of a lack of other abundant known U(VI)-reducing microorganisms. Monitoring the levels of methyl CoM reductase subunit A (*mcrA*) transcripts during an acetate-injection field experiment demonstrated that acetoclastic methanogens from the genus *Methanosarcina* were enriched after 40 days of acetate amendment. The increased abundance of *Methanosarcina* corresponded with an accumulation of methane in the groundwater. An enrichment culture dominated by a *Methanosarcina* species with the same *Methanosarcina mcrA* sequence that predominated in the field experiment could effectively convert acetate to methane. In order to determine whether *Methanosarcina* species could be participating in U(VI) reduction in the subsurface, cell suspensions of *M. barkeri* were incubated in the presence of U(VI) with acetate provided as the electron donor. U(VI) was reduced by metabolically active *M. barkeri* cells, however, no U(VI) reduction was observed in inactive controls. These results demonstrate that *Methanosarcina* species could play an important role in the long-term bioremediation of uranium-contaminated aquifers after depletion of Fe(III) oxides limits the growth of *Geobacter* species. The results also suggest that *Methanosarcina* have the potential to influence uranium geochemistry in a diversity of anaerobic sedimentary environments.

## Introduction

Injection of acetate into the groundwater of uranium-contaminated aquifers has been shown to be an effective way to stimulate microbially-mediated reductive precipitation of soluble U(VI) to poorly soluble U(IV) (Anderson *et al.*, 2003; Williams *et al.*, 2011; Williams *et al.*, 2013). A wide diversity of microorganisms are capable of U(VI) reduction (Kashefi & Lovley, 2000; Lovley *et al.*, 1991; Lovley *et al.*, 1993; Tebo & Obraztsova, 1998; Wall & Krumholz, 2006; Wu *et al.*, 2006) but only *Geobacter* species have been shown to reduce U(VI) with acetate as an electron donor. Although growth with acetate as the electron donor and U(VI) as the electron acceptor is possible (Lovley *et al.*, 1991), the low concentrations of U(VI), even in heavily contaminated subsurface environments requires that microbes use other forms of respiration as their primary means of energy conservation (Finneran *et al.*, 2002). *Geobacter* species grow rapidly in the initial phases of subsurface uranium bioremediation with added acetate because Fe(III) oxides are typically abundant in subsurface environments (Anderson *et al.*, 2003; Holmes *et al.*, 2002; Holmes *et al.*, 2005; Holmes *et al.*, 2007; Holmes *et al.*, 2013a) and *Geobacter* species outcompete other Fe(III) reducers under conditions of high acetate availability (Lovley *et al.*, 2011; Zhuang *et al.*, 2011). However, the potential for other microorganisms to contribute to acetate oxidation coupled to U(VI) reduction, especially after the Fe(III) oxides that support *Geobacter* growth are depleted, has not been intensively investigated. Sulfate reducers that can reduce U(VI) have been identified, but none of these are known to use acetate as an electron donor (Lovley & Phillips, 1992; Lovley *et al.*, 1993; Tebo & Obraztsova, 1998; Wall & Krumholz, 2006). Furthermore, relying on sulfate reducers to reduce U(VI) may not be a good long-term strategy because acetate additions can rapidly deplete sulfate from groundwater (Miletto *et al.*, 2011; N'Guessan *et al.*, 2008; Vrionis *et al.*, 2005).

Unlike Fe(III)- and sulfate-reducers, methanogens can thrive for long periods of time in organic-rich environments without external inputs of electron acceptors because they can conserve energy either from acetate dismutation or from the reduction of carbon dioxide, an electron acceptor generated by fermentation in their environment. If methanogens were capable of U(VI) reduction then this would make long-term *in situ* bioremediation of U(VI) a more attractive practice. To our knowledge, U(VI) reduction by methanogens has not been previously described. Previous studies have shown that methanogens can transfer electrons to various Fe (III) forms (Bodegom *et al.*, 2004; Bond & Lovley, 2002; Liu *et al.*, 2011; Sivan *et al.*, 2016; Vargas *et al.*, 1998; Zhang *et al.*, 2012), as well as vanadate (Zhang *et al.*, 2014) and humic substances (Cervantes *et al.*, 2002). However, acetate has not been shown to serve as an electron donor for these processes.

Evidence for methane production in response to acetate amendments during *in situ* uranium bioremediation (Holmes *et al.*, 2014) led us to investigate the potential for methanogens to further contribute to uranium bioremediation. The results suggest that *Methanosarcina* species that can couple the oxidation of acetate to the reduction of U(VI) might aid in the bioremediation process.

## Materials and methods

### Description of sampling site

The Rifle 24-acre experimental site is located close to the Colorado River, on the premises of an earlier uranium ore processing facility. Uranium concentrations in the water table of the Rifle aquifer are 2-8 times higher than the drinking water contamination limit (0.126 *μ*M) established by the uranium mill tailings remedial action (UMTRA). A detailed review of geochemical characteristics of the site has already been published (Zachara *et al.*, 2013) and *in situ* bioremediation of U^6+^ has been intensely studied at this site (Anderson *et al.*, 2003; Williams *et al.*, 2011; Williams *et al.*, 2013). Similar to previous years, acetate was injected into the subsurface at a concentration of ~15 mM between August and October, 2011 and monitored from 6 different wells (Giloteaux *et al.*, 2013). Groundwater and sediments for this study were collected from well CD-01 (a down gradient well) and a background well (CU-01) that never received any acetate additions.

### Enrichment of the dominant acetoclastic methanogens related to *Methanosarcina*

Methanogenic incubations were prepared by addition of 5 g wet sediment and 5 ml aquifer groundwater to 40 mL freshwater DSMZ 120 medium with acetate (40 mM) and no yeast extract in 160 mL serum bottles in an anaerobic chamber under an 80:20 N_2_:CO_2_ atmosphere. To reduce growth of bacteria, the antibiotics kanamycin (200 *μ*g/ml), erythromycin (200 *μ*g/ml), and penicillin-G (50 *μ*g/ml) were added to the slurries. All slurries were incubated for 30 days at 18°C in the dark.

In order to enrich for *Methanosarcina*, serial dilutions to extinction were carried out in 9 mL modified DSM 120 medium with acetate (40 mM). Methane production rates were determined after 8 transfers on DSM 120 medium. Phase contrast microscopy was performed on an Axioplan epifluorescence microscope on untreated cells enriched from Rifle sediment and aquifer water.

### Nucleic acid extraction and cDNA preparation

For nucleic acid extraction, it was first necessary to concentrate 50 L of groundwater by impact filtration on 293 mm diameter Supor membrane disc filters with pore sizes of 1.2 and 0.2 *μ*m (Pall Life Sciences). All filters were placed into whirl-pack bags, flash frozen in a dry ice/ethanol bath, and shipped on dry ice back to the laboratory where they were stored at −80°C. RNA was extracted from the filters using a modified phenol-chloroform method, as previously described (Holmes *et al.*, 2005). DNA was extracted from the filters with the FastDNA SPIN Kit for Soil (MP Biomedicals, Santa Ana, CA) according to the manufacturer’s instructions.

Extracted RNA and DNA were quantified with a NanoDrop spectrophotometer (Thermo Scientific, Wilmington, DE, USA) and stored at −80°C until further analyses. A DuraScript enhanced avian RT single-strand synthesis kit (Sigma, Sigma-Aldrich, St Louis, MO, USA) was used to generate cDNA from RNA, as previously described (Giloteaux *et al.*, 2013).

### PCR amplification parameters and microbial community analysis

For clone library construction, fragments from the *mcrA* gene which codes for the large subunit of methyl CoM reductase and the 16S rRNA gene were amplified from cDNA with mcrAf/mcrAr primers (Luton *et al.*, 2002) and with 344f/915r (Casamayor *et al.*, 2002) (Supplementary Table S1). Amplicons were ligated into the pCR-TOPO2.1 TA cloning vector according to manufacturer’s instructions (Invitrogen, the Netherlands). Inserts from the recombinant clones were directly amplified by PCR with M13 primers, purified and sequenced at the University of Massachusetts sequencing facility.

T-RFLP analysis was performed as described before (Shrestha et al. 2008). Gel-purified 5’-carboxyfluorescein-labeled 16S rRNA gene products amplified with Ar109f and Ar1000r were digested with TaqI at 37°C for 3 hours. The length of fluorescently labeled T-RFs was determined by comparison with the internal standard (LIZ 1200, ABI) using Genescan software (Applied Biosystems). The relative abundance of individual T-RFs in a given T-RFLP pattern was determined as the peak height of the respective T-RF divided by the total peak height of all T-RFs detected and was expressed as percentages (Dunbar et al., 2001; Shrestha et al., 2010). A clone library was also assembled from T-RFLP PCR products and 15 randomly selected clones were sequenced.

### Quantification of *Methanosarcina* mcrA transcript abundance

The quantitative PCR primer set (msa_mcrA173f/271r) designed to target *mcrA* genes from *Methanosarcina* species found in the Rifle subsurface was designed according to the manufacturer’s specifications (Applied Biosystems) (Supplementary Table S1). Quantitative PCR amplification and detection was performed with the 7500 Real Time System (Applied Biosystems) using cDNA made by reverse transcription from mRNA extracted from groundwater collected during the bioremediation experiment. Each reaction mixture consisted of a total volume of 25 *μ*l and contained 1.5 *μ*l of the appropriate primers (stock concentration 1.5 *μ*M), 5 ng cDNA, and 12.5 *μ*l Power SYBR Green PCR Master Mix (Applied Biosystems). All qPCR experiments followed MIQE guidelines (Bustin *et al.*, 2009) and qPCR efficiencies were 98%. Optimal thermal cycling parameters consisted of an activation step at 50°C for 2 minutes, an initial 10 minute denaturation step at 95°C, and 50 cycles of 95°C for 15 sec and 60°C for 1 minute. A dissociation curve generated by increasing the temperature from 58 to 95°C at a ramp rate of 2% showed that the PCR amplification process yielded a single predominant peak, further supporting the specificity of the qPCR primer pair.

### Phylogenetic analysis

16S rRNA and *mcrA* gene sequences were compared to Genbank nucleotide and protein databases with the BLASTn and BLASTx algorithms (Altschul *et al.*, 1990; Altschul *et al.*, 1997). Alignments were generated with MAFFT (Katoh & Standley, 2013) and PRANK (Loytynoja & Goldman, 2005) algorithms. The phylogenetic tree was inferred using the Neighbor Joining Method (Saitou & Nei, 1987). The percentage of replicate trees in which the associated taxa clustered together in the bootstrap test (100 replicates) is shown next to the branches (Felsenstein, 1985). All positions with less than 95% coverage were eliminated and a total of 93 positions were considered in the final dataset. All evolutionary analyses were conducted in MEGA7 (Kumar *et al.*, 2016).

Nucleotide sequences of *mcrA* genes used for phylogenetic analyses have been deposited in the Genbank database under accession numbers MF616623-MF616647.

### U(VI) reduction studies

Batch cultures of 500 mL *Methanosarcina barkeri* (DSM 800) were grown under strict anaerobic conditions (Balch *et al.*, 1979) on modified DSMZ medium 120 (Rotaru *et al.*, 2014) with acetate (40 mM) as substrate, and incubated at 37°C for ~3 weeks. Cultures were harvested when they reached an optical density at 600 nm of 0.19. All cell suspension preparations were performed in an anaerobic chamber to minimize oxygen exposure. Cells were pelleted by centrifugation for 10 minutes at 4000 x g in a Sorval RC 5B Plus centrifuge. These pellets were then washed twice in anoxic phosphate depleted buffer (PDB), which consisted of the following salts: 0.2 g/L MgSO_4_ × 7H_2_O, 0.025 g/L CaCl_2_ × 2H_2_O, 1 g/L NaCl, and 2 g/L NaHCO_3_. Cell pellets were then resuspended in 10 mL anoxic PDB to a cell density of ~0.4-0.5 at 600 nm. To generate heat-killed cells, 3 mL of this suspension was autoclaved at 122°C for 30 minutes. Six replicates were prepared by diluting 1 mL of the cell-suspension in 9 mL PDB buffer. For the heat-killed incubation, 1 mL autoclaved cell suspension was diluted in 9 mL PDB buffer. Sulfide (0.5 mM) was added to all inoculated tubes to ensure anoxic conditions. Acetate (40 mM) was also added to the tubes to fuel methanogenic activity. Triplicate live cell suspensions (active cells) and triplicate heat-killed controls (heat-killed cells) were incubated at 37°C. The other three live cell suspensions were incubated at 4°C (inactive cells). All cell suspensions were incubated with 0.2 mM U^6+^ prepared from a stock of uranyl-acetate (5 mM). Cell densities were determined with a bench top spectrophotometer, by absorbance measurements at 600 nm with mili-Q water as a blank.

The ability of *Methanosarcina barkeri* to reduce U(VI) was verified with U(VI) depletion measurements carried out on different cell suspensions over the course of 24 hours. Samples (0.1 mL) were retrieved anaerobically and diluted in 14.9 mL anoxic bicarbonate (100 mM) and 14.9 mL Uraplex solution. Concentrations of U(VI) were then measured with a kinetic phosphorescence analyzer, as previously described (Orellana *et al.*, 2013).

### Chemical analyses

Groundwater samples for geochemical analyses were collected after purging 12 L of groundwater from the wells with a peristaltic pump. The phenanthroline method was used to determine ferrous iron concentrations. Sulfate, and thiosulfate concentrations were measured with an ion chromatograph (ICS-2100, Dionex, CA) equipped with an AS18 column under isocratic elution with 32 mM KOH as the eluent. Acetate concentrations were determined with a high performance liquid chromatograph equipped with an ion exclusion HPX-87H column (Biorad, Hercules, CA) using 8 mM sulfuric acid as eluent. *In situ* methane production was monitored as previously described (Holmes *et al.*, 2014). Methane in the headspace of sediment/groundwater incubations was measured as previously described (Rotaru *et al.*, 2014) using a gas chromatograph with a flame ionization detector (Shimadzu, GC-8A).

## Results and discussion

### Evidence for acetoclastic methanogenic activity during acetate amendments

Methanogens that utilize acetate are restricted to the order Methanosarcinales (Kendal and Boone 2006). In order to determine whether the addition of acetate could promote the growth of acetoclastic methanogens in a uranium-contaminated aquifer, the activity of Methanosarcinales was investigated by monitoring *Methanosarcina mcrA* gene transcript abundance. Before day 39, less than 1.2 × 10^3^ *Methanosarcina mcrA* mRNA transcripts were detected per *μ*g of RNA extracted from the groundwater (Figure 1A). However, by day 46, *Methanosarcina mcrA* transcripts increased by 4 orders of magnitude to 3.7 × 10^7^ transcripts per *μ*g RNA. This increase in *Methanosarcina* coincided with a steep decline in groundwater sulfate concentrations (Figure 1B). Although sulfate reducers and methanogens compete for acetate (Lovley & Klug, 1983; Oremland & Polcin, 1982), high concentrations of acetate in the groundwater (Figure 1c) made it unlikely that growth of Methanosarcinales in the subsurface was being restricted by competition for acetate.

**Figure 1.**
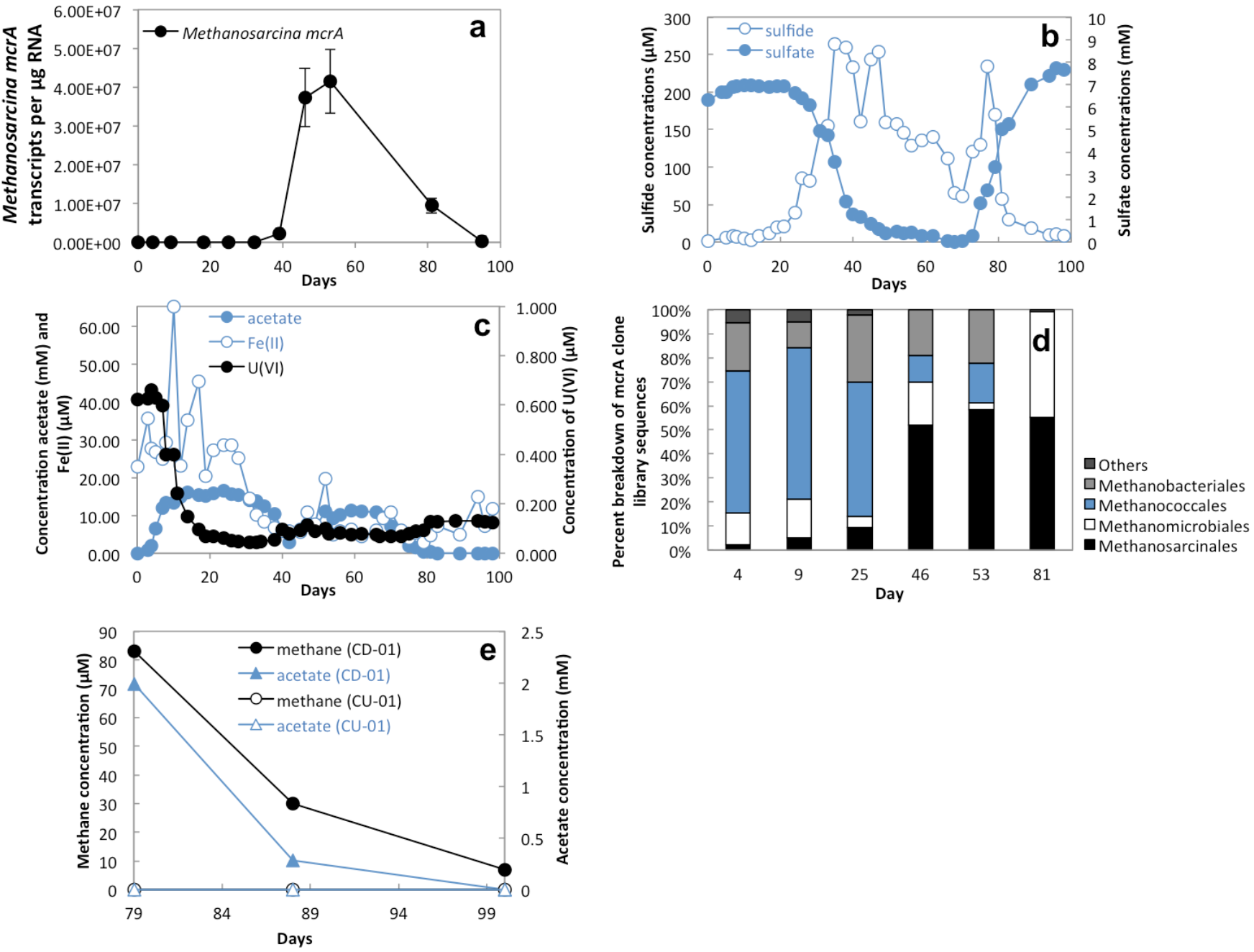
The injection of acetate into a uranium-contaminated aquifer, triggered acetate utilization coupled with iron reduction, sulfate reduction, and methanogenesis. (a) Quantitative RT-PCR of *Methanosarcina mcrA* mRNA transcripts recovered in the groundwater over the course of 100 days. (b) Concentrations of hydrogen sulfide (*μ*M) and sulfate (mM) detected in the groundwater. (c) Concentrations of acetate (mM), Fe(II) (*μ*M), and U(VI) (*μ*M) detected in the groundwater. (d) Proportion of *mcrA* sequences from various methanogenic families found in cDNA clone libraries assembled from RNA extracted from groundwater at different points during the experiment. (e) Concentrations of methane and acetate in an active well (CD-01) and a background well (CU-01) on days 79, 89, and 100. For further reference to geochemical parameters and cDNA clone libraries see Holmes et al. 2014.

The increase in Methanosarcinales coincided with an increase in free sulfide in the groundwater, producing highly reducing conditions that favor the growth of methanogens. Another consideration is the slow growth rate of Methanosarcinales, which might have limited their growth after acetate injections even under the most favorable conditions. The lack of sufficient reducing conditions coupled with the slow growth rate of Methanosarcinales may explain the finding that although acetate concentrations were high during the Fe(III) reducing phase of the experiment (days 0-33) (Figure 1c), the number of Methanosarcinales sequences stayed low until sulfate reduction became the primary subsurface metabolism (Figure 1a and 1d). The increase in abundance of Methanosarcinales was later followed by a decline, which coincided with acetate limitation associated with the halt in acetate injections on day 68.

Measurements of methane concentrations in the groundwater were not initiated until day 79 (Figure 1e). The high concentration of methane at this time demonstrated that methanogens had been active in the preceding days. Methane concentrations steeply declined over time coincident with the steep decline in acetate availability.

### Enrichment of *Methanosarcina* from Rifle

To determine whether *Methanosarcina* sequences detected in the groundwater came from metabolically active acetoclastic methanogens, methanogenic enrichments were established with groundwater and sediment collected from the uranium-contaminated site as inoculate and acetate (40 mM) as the substrate for growth. After 8 transfers, the enrichment cultures produced 96.04 *μ*moles/day^−1^ of methane, and the majority of cells were round, forming typical *Methanosarcina* rosettes (Fig. 2a). TRFLP analysis and sequencing of 16S rRNA gene fragments amplified from the enrichment cultures confirmed that the dominant TRF corresponded with *Methanosarcina* (Supplementary Figure S1).

**Fig. 2.**
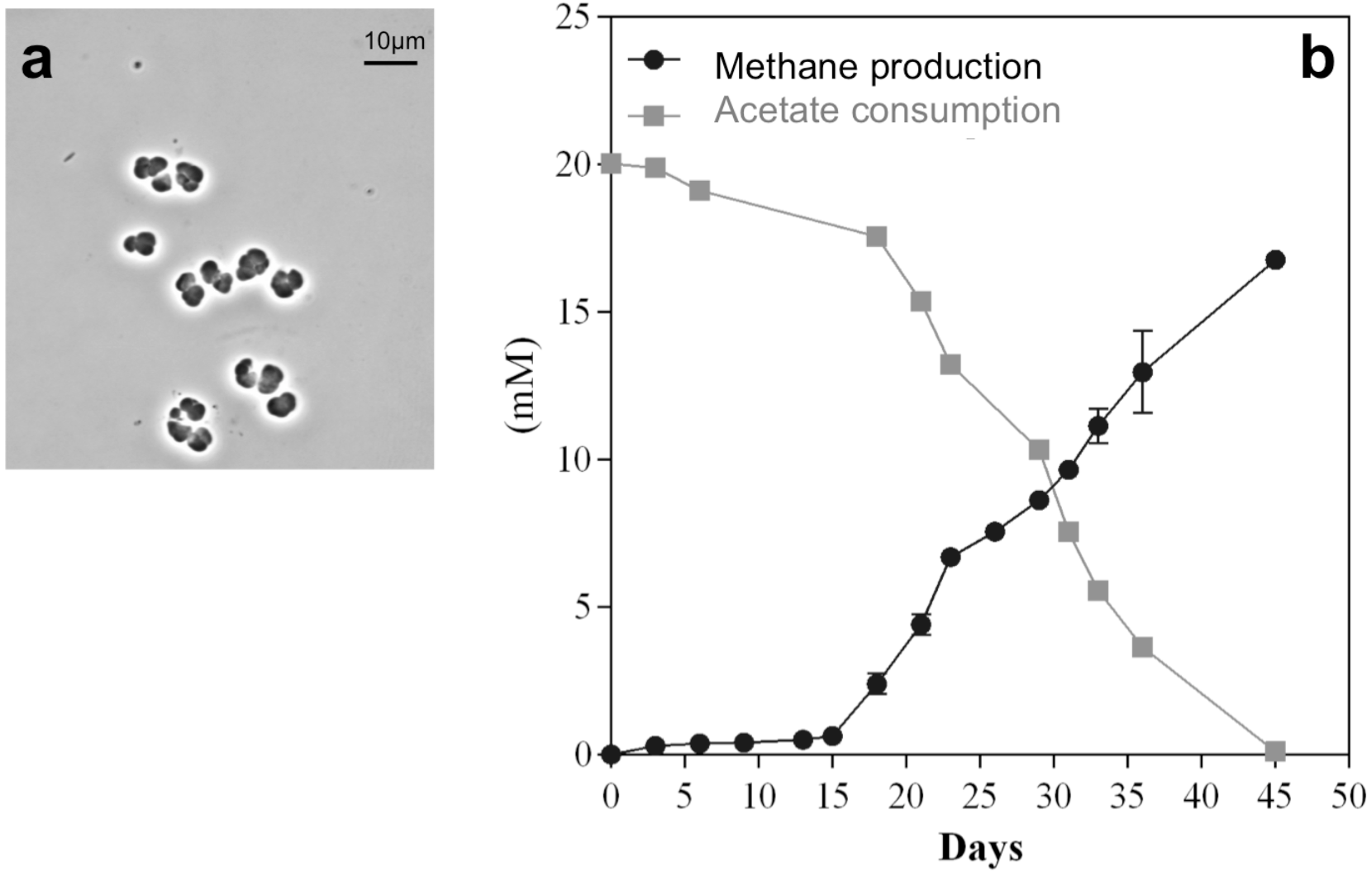
*Methanosarcina* enrichments from Rifle were acetoclastic. *Methanosarcina*-like cells enriched on 40 mM acetate from Rifle groundwater and sediment. Rifle *Methanosarcina* grew as rosettes (a) while consuming acetate and producing methane (b).

The dominant *Methanosarcina mcrA* cDNA sequence in the enrichment culture (Enrichment sequence 1 in Supplementary Figure S1B) was identical to the most abundant (48.2%) *Methanosarcina mcrA* cDNA sequence recovered from groundwater on day 53. Other *Methanosarcina mcrA* cDNA sequences detected on day 53 included sequences most similar to *M. barkeri* (37% of the sequences), *M. mazei* (11.1% of the sequences), and *M. acetivorans* (3.7% of the sequences).

This enrichment culture dominated by the most abundant *Methanosarcina* from the field experiment, could not grow with H_2_ or formate as electron donors, but grew on acetate at rates (Figure 2b) consistent with the emergence of *Methanosarcina* after approximately 40 days in the field experiment.

### U(VI) reduction by metabolically active *Methanosarcina* cells

To evaluate whether *Methanosarcina* species might be capable of U(VI) reduction, cell suspensions of *M. barkeri* were incubated with acetate as the electron donor and 200 *μ*M U(VI) as a potential electron acceptor. Within one day, the cells produced 1.6 mM methane while depleting 51% of the provided U(VI) (Fig. 3a). In contrast, cell suspensions incubated at 4 °C or autoclaved prior to incubation, did not produce methane or remove U(VI) (Figs 3b and 3c). These results indicated that U(VI) removal could be attributed to U(VI) reduction by metabolically active cells.

**Fig 3.**
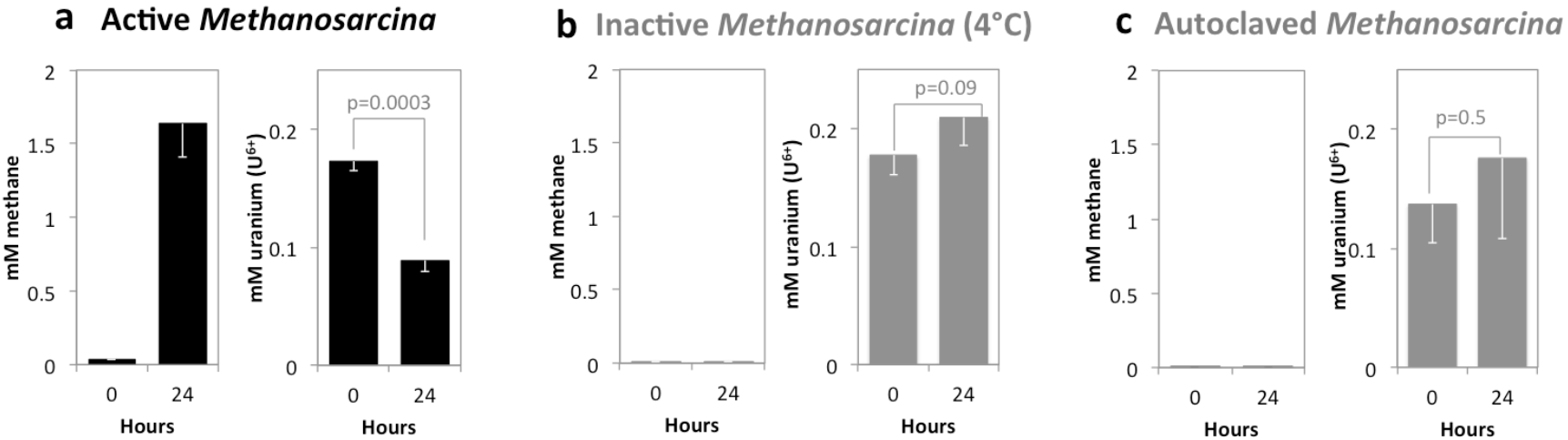
Uranium U(VI) reduction by metabolically active Methanosarcina cells. Metabolically active cells which were defined as such because they were producing methane from acetate were able to convert 51% of U(VI) to U(IV) (a) whereas metabolically inactive cells kept at 4°C in the same medium did not produce methane and also did not convert U(VI) to U(IV) (b), and neither did autoclaved cell suspensions from the same culture (c). The difference between original concentrations of U(VI) and the amount recovered in metabolically active cell suspensions after 24 hours of exposure was statistically different (p=0.0003).

### Implications

Our findings that acetate additions during *in situ* uranium bioremediation promotes the growth of *Methanosarcina* and that a *Methanosarcina* can reduce U(VI) has important implications for the design of long-term *in situ* uranium bioremediation strategies. Previous interpretations of U(VI) reduction during acetate-amendment at the Rifle, Colorado study site have focused on the U(VI) reduction capacity of *Geobacter* species because of their prevalence at the site (Anderson *et al.*, 2003; Wilkins *et al.*, 2009; Wilkins *et al.*, 2011; Wilkins *et al.*, 2013; Williams *et al.*, 2011; Williams *et al.*, 2013; Yun *et al.*, 2011) and because the sulfate-reducers that are enriched with acetate amendments (Dar *et al.*, 2013; Holmes *et al.*, 2013b; Miletto *et al.*, 2011; Vrionis *et al.*, 2005) are not likely to be effective U(VI) reducers. In fact, there has yet to be a description of an acetate-utilizing sulfate-reducing microorganism capable of U(VI) reduction. The results presented here suggest that *Methanosarcina* may also contribute to U(VI) reduction in the field experiments. Unlike *Geobacter* species, *Methanosarcina* do not require an external electron acceptor for acetate metabolism. Therefore, in longterm *in situ* uranium bioremediation *Methanosarcina* may emerge as an important microbial catalyst for uranium removal.

Furthermore, microbial reduction of U(VI) may play an important role in the uranium geochemistry of a diversity of sedimentary environments (Lovley *et al.*, 1991). Thus, the potential contribution of *Methanosarcina* to U(VI) reduction in anaerobic environments should be considered.

## Aknowledgements

This work is a contribution to grant: by the Office of Science (BER), US Department of Energy, Awards DE-SC0004080 and DE-SC0004814 and Cooperative Agreement DE-FC02-02ER63446. AER has been supported by grants from the Novo Nordisk Foundation, Innovationsfonden (410600017) and Danish Research Council (418100203). We would like to thank Devesh Shrestha for help with TRFLP experiments and Dan Carlo Flores for help starting enrichments.

## References

Altschul, S. F., Gish, W., Miller, W., Myers, E. W. & Lipman, D. J. (1990). Basic Local Alignment Search Tool. Journal of Molecular Biology 215, 403–410.

Altschul, S. F., Madden, T. L., Schaffer, A. A., Zhang, J., Zhang, Z., Miller, W. & Lipman, D. J. (1997). Gapped BLAST and PSI-BLAST: a new generation of protein database search programs. Nucleic Acids Res 25, 3389–3402.

Anderson, R. T., Vrionis, H. A., Ortiz-Bernad, I. & other authors (2003). Stimulating the in situ activity of Geobacter species to remove uranium from the groundwater of a uranium-contaminated aquifer. Appl Environ Microbiol 69, 5884–5891.

Balch, W. E., Fox, G. E., Magrum, L. J., Woese, C. R. & Wolfe, R. S. (1979). Methanogens: reevaluation of a unique biological group. Microbiol Rev 43, 260–296.

Bodegom, P. M., Scholten, J. C. & Stams, A. J. (2004). Direct inhibition of methanogenesis by ferric iron. FEMS Microbiol Ecol 49, 261–268

Bond, D. R. & Lovley, D. R. (2002). Reduction of Fe(III) oxide by methanogens in the presence and absence of extracellular quinones. Environmental Microbiology 4, 115–124.

Bustin, S. A., Benes, V., Garson, J. A. & other authors (2009). The MIQE Guidelines: Minimum Information for Publication of Quantitative Real-Time PCR Experiments. Clin Chem 55, 611–622.

Casamayor, E. O., Massana, R., Benlloch, S. & other authors (2002). Changes in archaeal, bacterial and eukaryal assemblages along a salinity gradient by comparison of genetic fingerprinting methods in a multipond solar saltern. Environ Microbiol 4, 338–348.

Cervantes, F. J., de Bok, F. A. M., Tuan, D. D., Stams, A. J. M., Lettinga, G. & Field, J. A. (2002). Reduction of humic substances by halorespiring, sulphate-reducing and methanogenic microorganisms. Environmental Microbiology 4, 51–57.

Dar, S., Tan, H., Peacock, A., Jaffe, P., N'Guessan, L., Williams, K. & Strycharz-Glaven, S. (2013). Spatial distribution of Geobacteraceae and sulfate-reducing bacteria during in situ bioremediation of uranium-contaminated groundwater. Remediation J 23, 31–49.

Dunbar, J., Ticknor, L., Kuske, C. (2001) Phylogenetic specificity and reproducibility and new method for analysis of terminal restriction fragment profiles of 16S rRNA genes from bacterial communities. Appl Environ Microbiol 67(1), 190–197

Felsenstein, J. (1985). Confidence-Limits on Phylogenies - an Approach Using the Bootstrap. Evolution 39, 783–791.

Finneran, K. T., Anderson, R. T., Nevin, K. P. & Lovley, D. R. (2002). Potential for Bioremediation of uranium-contaminated aquifers with microbial U(VI) reduction. Soil Sediment Contam 11, 339–357.

Giloteaux, L., Holmes, D. E., Williams, K. H. & other authors (2013). Characterization and transcription of arsenic respiration and resistance genes during in situ uranium bioremediation. ISME J 7, 370–383.

Holmes, D. E., Finneran, K. T., O'Neil, R. A. & Lovley, D. R. (2002). Enrichment of members of the family Geobacteraceae associated with stimulation of dissimilatory metal reduction in uranium-contaminated aquifer sediments. Appl Environ Microbiol 68, 2300–2306.

Holmes, D. E., Nevin, K. P., O'Neil, R. A., Ward, J. E., Adams, L. A., Woodard, T. L., Vrionis, H. A. & Lovley, D. R. (2005). Potential for quantifying expression of the Geobacteraceae citrate synthase gene to assess the activity of Geobacteraceae in the subsurface and on current-harvesting electrodes. Appl Environ Microbiol 71, 6870–6877.

Holmes, D. E., O'Neil, R. A., Vrionis, H. A. & other authors (2007). Subsurface clade of Geobacteraceae that predominates in a diversity of Fe(III)-reducing subsurface environments. Isme Journal 1, 663–677.

Holmes, D. E., Giloteaux, L., Barlett, M., Chavan, M. A., Smith, J. A., Williams, K. H., Wilkins, M., Long, P. & Lovley, D. R. (2013a). Molecular analysis of the in situ growth rates of subsurface Geobacter species. Appl Environ Microbiol 79, 1646–1653.

Holmes, D. E., Giloteaux, L., Williams, K. H., Wrighton, K. C., Wilkins, M. J., Thompson, C. A., Roper, T. J., Long, P. E. & Lovley, D. R. (2013b). Enrichment of specific protozoan populations during in situ bioremediation of uranium-contaminated groundwater. ISME J 7, 1286–1298.

Holmes, D. E., Giloteaux, L., Orellana, R., Williams, K. H., Robbins, M. J. & Lovley, D. R. (2014). Methane production from protozoan endosymbionts following stimulation of microbial metabolism within subsurface sediments. Front Microbiol 5.

Kashefi, K. & Lovley, D. R. (2000). Reduction of Fe(III), Mn(IV), and toxic metals at 100 degrees C by Pyrobaculum islandicum. Appl Environ Microb 66, 1050–1056.

Katoh, K. & Standley, D. M. (2013). MAFFT Multiple Sequence Alignment Software Version 7: Improvements in Performance and Usability. Mol Biol Evol 30, 772–780.

Kendal, M., Boone, D. (2006). The order Methanosarcinales. In The Prokaryotes. pp. 244–256. Edited by Dworkin, M.

Kumar, S., Stecher, G. & Tamura, K. (2016). MEGA7: Molecular Evolutionary Genetics Analysis Version 7.0 for Bigger Datasets. Mol Biol Evol 33, 1870–1874.

Liu, D., Dong, H. L., Bishop, M. E., Wang, H. M., Agrawal, A., Tritschler, S., Eberl, D. D. & Xie, S. C. (2011). Reduction of structural Fe(III) in nontronite by methanogen Methanosarcina barkeri. Geochim Cosmochim Ac 75, 1057–1071.

Lovley, D. R. & Klug, M. J. (1983). Sulfate Reducers Can out-Compete Methanogens at Fresh-Water Sulfate Concentrations. Appl Environ Microbiol 45, 187–192.

Lovley, D. R., Phillips, E. J. P., Gorby, Y. A. & Landa, E. R. (1991). Microbial Reduction of Uranium. Nature 350, 413–416.

Lovley, D. R. & Phillips, E. J. P. (1992). Bioremediation of Uranium Contamination with Enzymatic Uranium Reduction. Environ Sci & Technol 26, 2228–2234.

Lovley, D. R., Roden, E. E., Phillips, E. J. P. & Woodward, J. C. (1993). Enzymatic Iron and Uranium Reduction by Sulfate-Reducing Bacteria. Marine Geology 113, 41–53.

Lovley, D. R., Ueki, T., Zhang, T. & other authors (2011). Geobacter: The Microbe Electric's Physiology, Ecology, and Practical Applications. In Adv Microb Phys, pp. 1-100. Edited by R. K. Poole.

Loytynoja, A. & Goldman, N. (2005). An algorithm for progressive multiple alignment of sequences with insertions. PNAS 102, 10557–10562.

Luton, P. E., Wayne, J. M., Sharp, R. J. & Riley, P. W. (2002). The mcrA gene as an alternative to 16S rRNA in the phylogenetic analysis of methanogen populations in landfill. Microbiology 148, 3521–3530.

Miletto, M., Williams, K. H., N'Guessan, A. L. & Lovley, D. R. (2011). Molecular Analysis of the Metabolic Rates of Discrete Subsurface Populations of Sulfate Reducers. Appl Environ Microbiol 77, 6502–6509.

N'Guessan, A. L., Vrionis, H. A., Resch, C. T., Long, P. E. & Lovley, D. R. (2008). Sustained removal of uranium from contaminated groundwater following stimulation of dissimilatory metal reduction. Environ Sci Technol 42, 2999–3004.

Orellana, R., Leavitt, J. J., Comolli, L. R. & other authors (2013). U(VI) Reduction by Diverse Outer Surface c-Type Cytochromes of Geobacter sulfurreducens. Appl Environ Microb 79, 6369–6374.

Oremland, R. S. & Polcin, S. (1982). Methanogenesis and sulfate reduction: competitive and noncompetitive substrates in estuarine sediments. Appl Environ Microbiol 44, 1270–1276.

Rotaru, A. E., Shrestha, P. M., Liu, F., Markovaite, B., Chen, S., Nevin, K. P. & Lovley, D. R. (2014). Direct interspecies electron transfer between Geobacter metallireducens and Methanosarcina barkeri. Appl Environ Microbiol 80, 4599–4605.

Saitou, N. & Nei, M. (1987). The neighbor-joining method: a new method for reconstructing phylogenetic trees. Mol Biol Evol 4, 406–425.

Sivan, O., Shusta, S. S. & Valentine, D. L. (2016). Methanogens rapidly transition from methane production to iron reduction. Geobiology 14, 190–203.

Shrestha, M., Abraham, W.-R., Shrestha, P. M., Noll, M., Conrad R. (2008). Activity and composition of methanotrophic bacterial communities in planted rice soil studied by flux measurements, analyses of pmoA gene and stable isotope probing of phospholipid fatty acids. Environ Microbiol 10 (2), 400–412.

Shrestha, M., Shrestha, P.M., Frenzel, P., Conrad, R. (2010). Effect of nitrogen fertilization on methane oxidation, abundance, community structure, and gene expression of methanotrophs in the rice rhizosphere. ISME J 4, 1545–1556.

Tebo, B. M. & Obraztsova, A. Y. (1998). Sulfate-reducing bacterium grows with Cr(VI), U(VI), Mn(IV), and Fe(III) as electron acceptors. Fems Microbiol Lett 162, 193–198.

Vargas, M., Kashefi, K., Blunt-Harris, E. L. & Lovley, D. R. (1998). Microbiological evidence for Fe(III) reduction on early Earth. Nature 395, 65–67.

Vrionis, H. A., Anderson, R. T., Ortiz-Bernad, I. & other authors (2005). Microbiological and geochemical heterogeneity in an in situ uranium bioremediation field site. Appl Environ Microbiol 71, 6308–6318.

Wall, J. D. & Krumholz, L. R. (2006). Uranium reduction. Annu Rev Microbiol 60, 149–166.

Wilkins, M. J., VerBerkmoes, N. C., Williams, K. H. & other authors (2009). Proteogenomic Monitoring of Geobacter Physiology during Stimulated Uranium Bioremediation. Appl Environ Microbiol 75, 6591–6599.

Wilkins, M. J., Callister, S. J., Miletto, M., Williams, K. H., Nicora, C. D., Lovley, D. R., Long, P. E. & Lipton, M. S. (2011). Development of a biomarker for Geobacter activity and strain composition; Proteogenomic analysis of the citrate synthase protein during bioremediation of U(VI). Microb Biotech 4, 55–63.

Wilkins, M. J., Wrighton, K. C., Nicora, C. D. & other authors (2013). Fluctuations in Species-Level Protein Expression Occur during Element and Nutrient Cycling in the Subsurface. PLOS ONE 8, e57819.

Williams, K. H., Long, P. E., Davis, J. A. & other authors (2011). Acetate Availability and its Influence on Sustainable Bioremediation of Uranium-Contaminated Groundwater. Geomicrobiol J 28, 519–539.

Williams, K. H., Bargar, J. R., Lloyd, J. R. & Lovley, D. R. (2013). Bioremediation of uranium-contaminated groundwater: a systems approach to subsurface biogeochemistry. Curr Opin Biotechnol 24, 489–497.

Wu, Q., Sanford, R. A. & Loffler, F. E. (2006). Uranium(VI) reduction by Anaeromyxobacter dehalogenans strain 2CP-C. Appl Environ Microb 72, 3608–3614.

Yun, J., Ueki, T., Miletto, M. & Lovley, D. R. (2011). Monitoring the Metabolic Status of Geobacter Species in Contaminated Groundwater by Quantifying Key Metabolic Proteins with Geobacter-Specific Antibodies. Appl Environ Microbiol 77, 4597–4602.

Zachara, J. M., Long, P. E., Bargar, J. & other authors (2013). Persistence of uranium groundwater plumes: Contrasting mechanisms at two DOE sites in the groundwater-river interaction zone. J Contam Hydrol 147, 45–72.

Zhang, J., Dong, H. L., Liu, D., Fischer, T. B., Wang, S. & Huang, L. Q. (2012). Microbial reduction of Fe(III) in illite-smectite minerals by methanogen Methanosarcina mazei. Chem Geol 292, 35–44.

Zhang, J., Dong, H. L., Zhao, L. D., McCarrick, R. & Agrawal, A. (2014). Microbial reduction and precipitation of vanadium by mesophilic and thermophilic methanogens. Chem Geol 370, 29–39.

Zhuang, K., Izallalen, M., Mouser, P., Richter, H., Risso, C., Mahadevan, R. & Lovley, D. R. (2011). Genome-scale dynamic modeling of the competition between Rhodoferax and Geobacter in anoxic subsurface environments. ISME J 5, 305–316.

